# Extracellular spike waveform analysis reveals cell type-specific changes in the superior colliculus of fragile X mice

**DOI:** 10.1101/2025.07.28.667270

**Authors:** Gourav Sharma, Ashley L. Russell, Karen G. Dixon, Romina Fusha, Jason W. Triplett

## Abstract

A long-standing goal of neuroscience has been to elucidate the diverse complement of neurons in the brain, which can be defined by several criteria. Analysis of action potential shape in extracellular recordings has revealed subpopulations in several regions of the brain, allowing for insights into neuronal subtype-specific function in the intact brain. The superior colliculus (SC) is a critical sensorimotor region, integrating visual, somatosensory, auditory, and nociceptive inputs to direct complex behaviors. Recent work suggests that the SC may be adversely impacted in neurodevelopmental disorders (NDDs), underscoring its importance. However, our understanding of cellular diversity in the SC lags in comparison to other regions, limiting our ability to parse circuit changes in NDDs. Here, we utilized semi-automated clustering methods to classify neurons in the mouse SC based on multiple features of extracellularly recorded waveforms to identify five putative cell types. Secondary analysis of firing statistics and visual tuning properties supported the cluster segregation. Interestingly, the proportions of units assigned to each cluster differed in the SC of a mouse model of fragile X syndrome (FXS, *Fmr1^-/y^*), with only four of five types identified. Furthermore, we observed changes in waveform properties and firing statistics, but not visual tuning properties, between genotypes in a subtype-specific manner. Taken together, these data add to our understanding of neuronal diversity in the SC and alterations of visual circuit organization and function in NDDs.

## Introduction

Understanding the diversity of neuronal cell types and how they contribute to brain function has been a central goal of neuroscience. Neuronal subtypes can be classified based on their morphology, connectivity, gene expression, function, and intrinsic electrophysiological properties, including the shape of the action potential waveform. *In vitro* studies initially showed differences in firing patterns and spike waveforms between cortical pyramidal neurons and GABAergic interneurons [1, 2]. Distinctions in extracellular waveform shape generally translate to intracellular features observed *in vivo* [3], allowing insights into subtype-specific contributions to network activity and responses to stimuli. However, analysis of only two broad categories does not reflect the rich neuronal diversity observed in many brain regions.

Recent evidence suggests that extracellular waveform analysis can be used to identify multiple subtypes of neurons [4-8]. While initial studies utilized only a few metrics derived from waveforms [2, 9-14], more recent approaches utilized a wider array of metrics to identify distinct shapes through a manual, hierarchical classification system [6, 7]. However, this approach may overlook nuances at the borders of metric thresholds and may have limited applicability to other brain regions where cutoffs would need to be determined empirically. In contrast, unbiased methods such as semi-supervised deep learning and machine learning have yielded better identification of neuronal subtypes [4, 5, 8]. However, these approaches are computationally intense and require large datasets to train and validate findings. While high-density recordings are becoming more prevalent, clustering methods that are both unbiased and applicable to smaller datasets could be broadly used.

The mouse superior colliculus (SC) has become an important model for investigating sensory circuit development, organization, and function [15, 16]. Classically understood to regulate head and eye movements [17], several recent lines of study suggest the SC plays a critical role in processing visual information to guide complex behaviors, including visual threat detection, predation, and decision making [18-21]. Thus, understanding its circuitry has become an active area of investigation. Morphologic and intrinsic electrophysiological studies suggest 4-6 distinct neuronal types in the rodent SC [22-24]. In contrast, functional and molecular studies suggest broader diversity may exist [25-29]. Whether distinct extracellular waveforms are discernible in the SC remains unknown.

Underscoring the need to interrogate circuit function in the SC is that it may be negatively impacted in neurodevelopmental disorders [30]. Fragile X syndrome (FXS) is caused by the silencing of the *FMR1* gene, which encodes fragile-X messenger ribonucleoprotein 1 (FMRP), and is the most prevalent single gene cause of autism spectrum disorder [31]. Previously, we reported disruptions in both anatomical organization and visual function in the SC of a mouse model of FXS (*Fmr1^-/y^*) [32]. Specifically, receptive fields were enlarged, direction-selectivity was reduced, axis-selective neurons were hyperactive, and inputs from layer 5 of V1 were disorganized in the SC of *Fmr1^-/y^* mice. However, it remains unclear how cell type diversity or subtype-specific function may be impacted in the FXS SC.

Here, we analyzed extracellular spike waveforms from visually responsive neurons in the SC of control (Fmr1^+/y^) and *Fmr1^-/y^* mice to determine if functionally distinct neuronal subtypes are identifiable and, if so, how they may be impacted by loss of FMRP expression. We used independent semi-automated clustering methods to identify five putative neuronal subtypes in the mouse SC based on metrics derived from individual waveforms. *Post hoc* analysis of firing statistics and visual tuning properties confirmed the subdivisions established by our clustering algorithm. Intriguingly, in *Fmr1^-/y^* mice, the proportions of neuronal subtypes were significantly altered, with one entire class absent. Furthermore, we observed changes in waveform properties and firing statistics and between genotypes in a subtype-specific manner. Collectively, these findings demonstrate the presence of diverse cell-types in the SC based on extracellular waveform shape and strongly suggest that FMRP is indispensable for the proper development of visual circuit organization and function within the SC.

## Materials and Methods

### Mice

*Fmr1* knockout (*Fmr1^-/y^*) mice were generated and genotyped as previously described [33]. Mice were maintained on a mixed C57BL/6 and 129/Sv background. Only male mice were used for both experimental and control animals to avoid potential mosaicism associated with heterozygous females [34, 35] and to facilitate the use of littermates as controls. Mice were housed in a temperature and humidity-controlled room under standard 12/12-h light-dark cycle. After weaning, mice were housed in groups of 1–5 with same-sex siblings. All procedures were performed in accordance with, and approved by, the Children’s National Research Institute’s IACUC.

### *In vivo* electrophysiology

Briefly, male mice (postnatal day 14 (P14)–P120) were anesthetized with isoflurane. The animal’s temperature was monitored and maintained at 37 °C through a feedback-controlled heating pad. Silicone oil was applied on the eyes to prevent drying. A craniotomy was performed on the right hemisphere ∼ 0.5 mm lateral from the midline suture and between 1.5 to − 0.25 mm anterior from the lambda suture.

Recordings were performed as previously described [32]. A 16-channel silicone probe (NeuroNexus) coated in 1,1’-Dioctadecyl-3,3,3’,3’-Tetramethylindocarbocyanine Perchlorate (DiI, ThermoFisher) was lowered between 0.8 and 1.5 mm into the SC at a 35° angle and stabilized with agarose (1% in PBS) (Fig. 1a & b). Electrical signals were amplified and filtered between 0.7 and 7 kHz, sampled at 25 kHz, and acquired using a System 3 workstation (Tucker-Davis Technologies).

**Figure 1:**
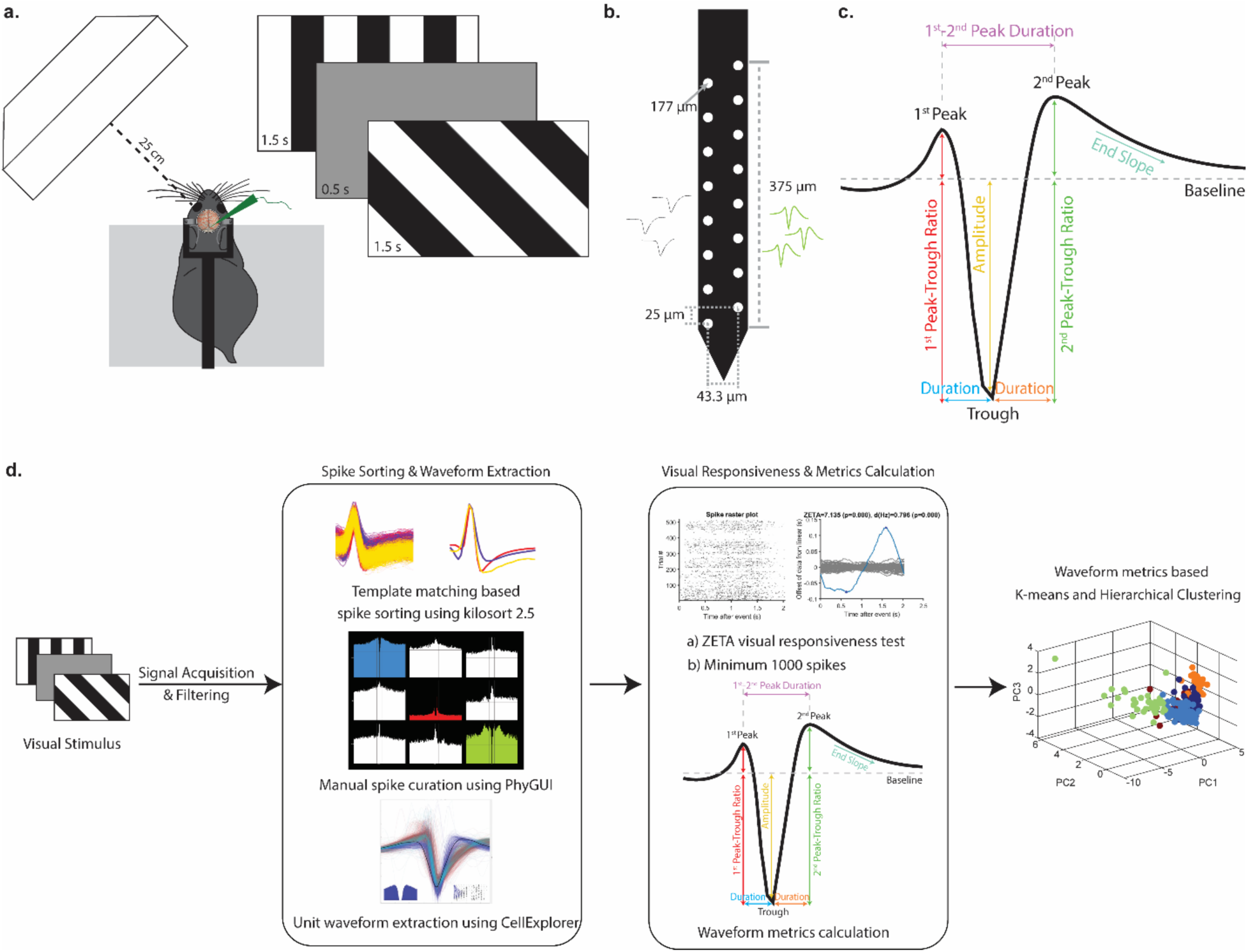
Schematics of the experimental and analysis methodologies. (a) Schematic of *in-vivo* visual presentation set up and square wave drifting grating paradigm. (b) Schematic of the silicon multi-electrode array utilized. (c) Example extracellular spike waveform with three major deflections (1^st^ Peak, Trough and 2^nd^ Peak) from baseline (*dashed line*). Spikes are classified based on six features: 1^st^ Peak-Trough Ratio (*red*); 2^nd^ Peak-Trough Ratio (*green*); 1^st^ Peak-2^nd^ Peak Duration (*purple*); 1^st^ Peak-Trough Time (*blue*); Trough-2^nd^ Peak Time (*orange*); and End Slope (*cyan*). (d) Schematic representation of the methodology used for classification of SUs and their waveform metrics-based clustering.

Immediately after *in vivo* recording mice were euthanized and intracardially perfused with ice-cold PBS followed by 4% paraformaldehyde (PFA). The brains were dissected out and post-fixed in 4% PFA overnight. The brains were then briefly washed in PBS and were embedded in 2% agarose and 50-μm sections were cut in the sagittal plane with a Manual Slice Vibratome (World Precision Instruments) in order to image the DiI-labeled probe penetrations in the SC. All imaging was performed on a BX63 microscope equipped with an AxioCam HR digital camera (Olympus).

### Visual stimuli

Visual stimuli were generated using custom software (MATLAB) using Psychtoolbox-3 [36]. The monitor (52 × 29.5 cm, 60-Hz refresh rate) was placed 25 cm from the animal in front of the eye contralateral to the recording penetration, subtending ∼ 90 × 70° of visual space (Fig. 1a). Drifting square waves of 100% contrast at 12 different orientations (30° spacing) and six different spatial frequencies between 0.01 and 0.32 cycles per degree (six logarithmic steps) were presented. A temporal frequency of 2 Hz was consistent for all the gratings. Each stimulus of given orientation and spatial frequency (or a blank condition) was presented for 1.5 s in a pseudorandom order for five trials, with an interstimulus interval of 0.5 s.

### Spike sorting

Signals were automatically processed via KiloSort 2.5 [37], a spike-sorting program designed for dense arrays to rectify the spatio-temporal issues from extracellular recordings. The outputs of KiloSort were then loaded into phy, an open-source visualization interface for manual curation [38]. First, clusters were ‘cleaned’ by manually drawing a boundary in the principal component analysis (PCA) space to remove abnormal spikes outside of the boundary. Next, an auto-correlogram of spike times and average spike waveforms were inspected to classify clusters as ‘noise’, ‘good’, or ‘multi-unit activity (MUA)’. Merges were made between ‘good’ clusters when. ‘Good’ clusters were merged if the cluster of spikes in the PCA overlapped and their cross-correlogram showed a >1 ms gap near zero or if there was evidence of drift when the clusters had similar waveforms, similar post-stimulus time histograms (PSTHs), similar auto-correlograms, and convergence in the amplitude versus time plot (i.e. one cluster stops spiking and the other starts at the same time). Next, ‘good’ clusters which had >1000 spikes across the recording session were identified as single units (SUs). Finally, SUs classified as visually responsive based on the parameter-free ZETA test [39] were chosen for further analysis.

### Visual tuning analysis

To calculate directional and axial tuning, we calculated a global orientation selectivity index (gOSI), which is the vector sum of response at each axis normalized by the scalar sum of responses:

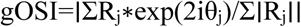

Where R_j_ is the neuronal response (e.g., spike rate) at orientation angle θ_j_ in radians presented at the preferred spatial frequency. A cell was considered axis selective (AS) with a gOSI greater than 0.2. For all AS neurons, the preferred orientation (tpref) was defined as that evoked the maximum spiking response across all spatial frequencies. Tuning curves were fitted with a sum of two Gaussians centered at q_pref_ and q_pref_ + p using the *nlinfit* function in MATLAB, and the tuning width was calculated as the half-width at half-maximum of the fitted curve. Linearity of spatial summation was calculated by applying a Fourier transform to binned spiking data from trials at q_pref_ and determining the ratio of the response at the first harmonic (F_1_) to the mean response (F_0_) [40]. Stimulus response latency was determined as the center of the first of 20 consecutive 20 msec sliding windows in which the firing rate was 2 SDs above or below the spontaneous rate at the neuron’s preferred orientation and spatial frequency.

### Waveform analysis

Average waveforms for SUs were extracted via Cell Explorer [41] and normalized to the absolute value of their peak deviation from baseline. For each waveform, the trough was identified using the *min* function in MATLAB and then a binomial curve fitted to the trough and adjacent two samples in both directions using the *polyfit* function. The first and second peaks were identified using the *findpeaks* function in the portion of the waveform before or after the trough, respectively, and again fitted with a binomial curve. To determine the end slope, a linear regression was fit through the sample occurring 0.5 msec after the trough and those immediately before and after this sample.

### Clustering methods

#### k-means clustering

As input for the clustering algorithm, we used the normalized spike waveform shape metrics that compared six features 1^st^ Peak-Trough Ratio, 2^nd^ Peak-Trough Ratio, 1^st^ Peak-2^nd^ Peak Duration, 1^st^ Peak-Trough Duration, Trough-2^nd^ Peak Time, and End-slope. Clustering was performed using the *kmeans* function in MATLAB, using the default settings for distance measure (squared Euclidean) and centroid seed locations (random data point), which yielded cluster assignments and distances to the centroid for each visually responsive unit. Silhouette coefficients were determined using the *silhouette* function in MATLAB, using the default settings for distance (squared Euclidean). We performed five independent iterations of *k*-means clustering with accompanying silhouette analysis.

#### Hierarchical clustering

To classify neuronal units by waveform characteristics, we performed hierarchical clustering in MATLAB. Input features included the same waveform metrics used for *k*-means clustering. All features were z-score normalized to ensure comparability. Pairwise Euclidean distances were computed, and clustering was performed using Ward’s linkage method, which minimizes within-cluster variance. An elbow-based cutoff determined the optimal number of clusters.

### Analysis of firing properties

Firing properties were quantified for each unit during the full trial window, which included both stimulus and inter-stimulus periods (excluding blank trials). For each trial, the firing rate was calculated as the number of spikes divided by the trial duration. To assess variability in spike counts across trails, the Fano factor was calculated as the ratio of the variance to the mean of trial firing rates. Temporal irregularity of spiking was measured using the coefficient of variation of inter-spike intervals (ISIs), calculated as the standard deviation of ISIs divided by the mean ISI. Finally, burstiness was evaluated using a burst index (BI), defined as the ratio of the observed proportion of ISIs ≤ 5ms to the proportion predicted by a Poisson distribution with the same mean firing rate.

### Code Availability

MATLAB codes used for clustering, determining visual tuning, and firing statistics are available at the Triplett Lab github: https://github.com/Triplett-Lab/Waveform2025.git.

### Statistical analysis

All values are reported as mean ± SEM. All data were first tested for normality of distribution using the D’Agostino & Pearson test. Levene’s test was used to evaluate equality of variances. Log transformations were applied to normalize skewed data and reduce variance heterogeneity where needed. For one-way comparisons, ordinary one-way ANOVA was used when variances were equal. When variances were unequal, we used Brown-Forsythe and Welch ANOVA. For lognormal data with equal variances, the Kruskal-Wallis test followed by Dunn’s multiple comparisons was used. Where appropriate, Tukey’s *post hoc* test was applied for equal variances, and Games-Howell for unequal variances. For two-way ANOVA, standard main-effect models were used to compare the effects of genotype and stimulus condition. Holm-Sidak *post hoc* tests were applied only for planned comparisons between specific groups. All tests were two-tailed, with significance set at p < 0.05. All analyses were performed with the statistical software package Prism (GraphPad).

## Results

### Segregation of multiple types of units in the SC based on spike waveform

We analyzed waveforms of visually responsive SUs identified in the SC and utilized semi-automated *k*-means clustering based on waveform properties (Fig. 1c). We calculated 6 metrics from normalized waveforms: 1^st^ peak-trough ratio, 1^st^-2^nd^ peak duration, 1^st^ peak-trough duration, through-2^nd^ peak duration, 2^nd^ peak-trough ratio and end-slope. From these data, we identified principal components and used those accounting for at least 10% of the variance for clustering via *k*-means (Fig. 2a).

**Figure 2:**
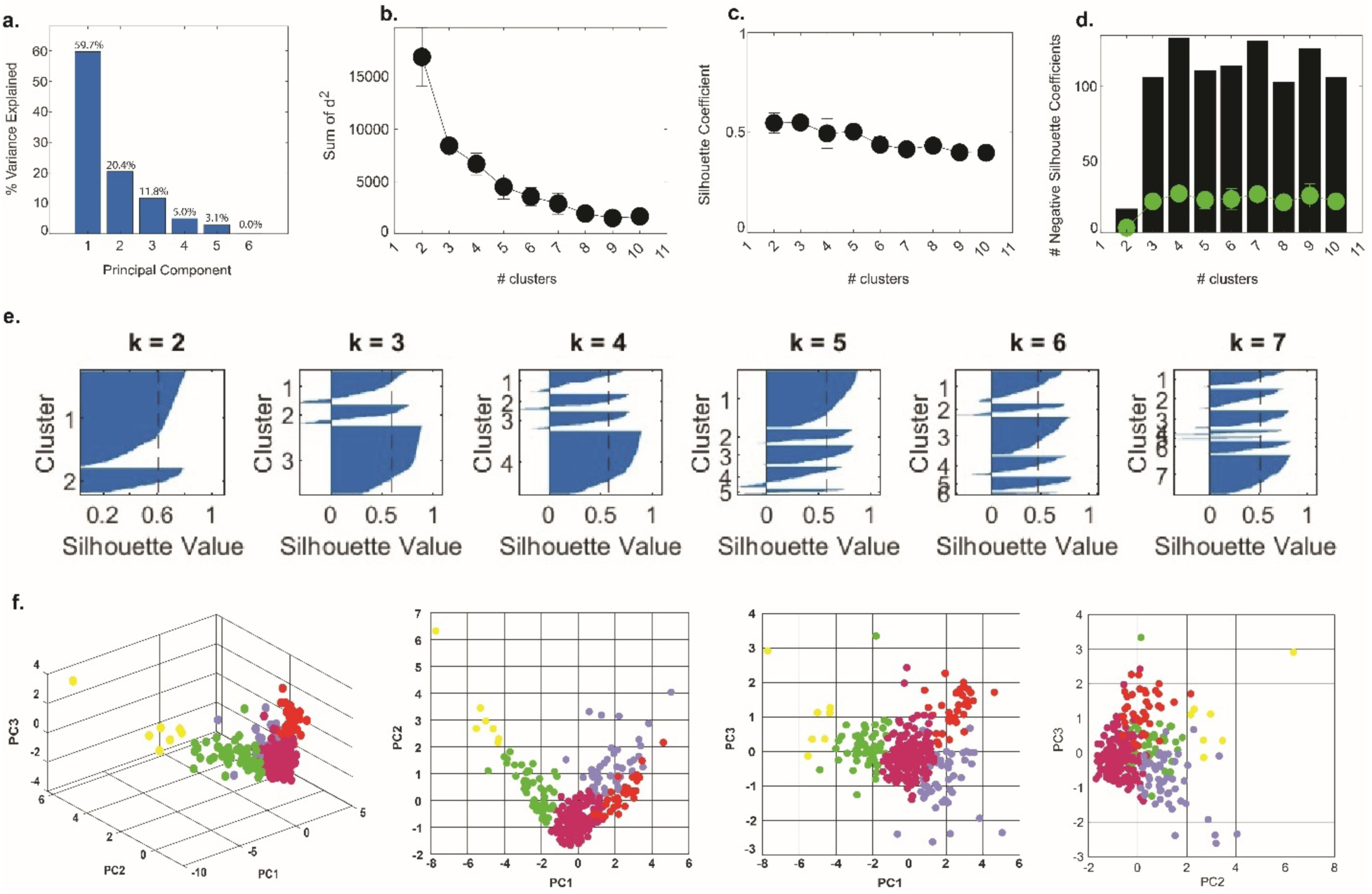
Semi-automated *k*-means clustering based on waveform features. (a) The contribution of each principal component to the explained variance across clusters. (b) Quantification of the sum of squared distances between units and assigned cluster centroid. (c) The mean silhouette coefficient across clusters. (d) The cumulative (black bars) and mean (green dots) number of negative silhouette values. (e) Representative plots of individual silhouette coefficients for units in their assigned cluster for the indicated number of clusters (“*k*”). (f) Distribution of identified clusters in the primary principal component spaces (PC1 vs PC2 vs PC3, PC1 vs PC2, PC1 vs PC3, PC2 vs PC3).

To determine the best value for *k* in our data set, we tested *k* values ranging from two through 10 and performed five iterations of clustering for each value. First, we examined the sum of squared distances between individual points and their centroid as a function of *k* (Fig. 2b). The resulting plot revealed that subdividing our data into more than five clusters does not substantially improve distance minimization. Next, we assessed the likelihood that points were assigned to the appropriate cluster by calculating the mean silhouette coefficient (Fig. 2c). We observed a decline in mean silhouette coefficient when *k* was greater than five, suggesting inappropriate cluster assignments. Analysis of the cumulative and mean number of negative silhouette coefficients revealed a reduction when the number of clusters was increased from four to five, suggesting slightly more appropriate clustering (Fig. 2d). Finally, we manually inspected silhouette plots for each value of *k* from a representative iteration (Fig. 2e). When *k* = 4 or 5, all clusters were comprised of a substantial number of members well above or very near the mean silhouette coefficient. Negative silhouette coefficients were observed in 3 of 4 clusters when *k* = 4, but only 1 of 5 clusters when *k* = 5, suggesting more accurate cluster assignments. To further validate this, we visualized clusters by plotting principal component values in 3D and 2D space (Fig. 2f & g). These plots revealed clearly separated clusters. Based on the relatively low sum of squared distances, high mean silhouette coefficient, low number of negative silhouette coefficients, robustness of each cluster relative to the mean silhouette coefficient, and clean separation in principal component space, we chose a value of *k* = 5 for further analysis.

A good separation of the clusters based on the waveform metrics should be visible as differences in the mean waveforms of each cluster. Indeed, we see clear differences in the mean waveforms of each of the clusters and good correspondence between individual units (Fig. 3a), supporting proper classification. We found that about half of the SUs (50.85%, 149/293) were classified into Cluster 5, which had a shape akin to narrow spiking neurons described in cortex (Fig. 3a, *purple*). The next most common subtype (Cluster 3, [17.75%, 52/293]), had a complex shape with a substantial positive peak prior to a larger, narrow trough and sharp downward end slope (Fig. 3a, *green*). Spikes in Cluster 1 (11.6%, 34/193) resemble classic broad/regular spiking neurons previously described (Fig. 3a, *red*). Intriguingly, Cluster 4, comprising 17.41% (51/293) of the population appeared to be intermediate between narrow and broad spiking, having no peak prior to a trough similar to broad spiking neurons and a negative end slope akin to narrow spiking neurons (Fig. 3a, *blue*). Finally, Cluster 2, which represented only 2.39% (7/293) of the population, had a complex waveform with a large initial peak, suggesting these may be axonal signals or back propagating dendritic signals (Fig. 3a, *yellow*) [42, 43]. To determine if waveforms in the SC could be similarly segregated based on specific waveform metrics as in previous studies in V1 [6, 7, 12], we plotted different combinations of metrics against one another (Fig. 3c-e). We observe that end slope plotted against the 1^st^ Peak-2^nd^ Peak duration shows the best separation amongst all the clusters (Fig. 3c). A few other combinations of metrics like end slope against Trough-2^nd^ peak duration (Fig. 3d) and end slope against 1^st^ Peak-Trough duration (Fig. 3e) also show clear separation of clusters. Taken together, these data suggest that five distinct waveform shapes can be identified in the visual layers of the mouse SC.

**Figure 3:**
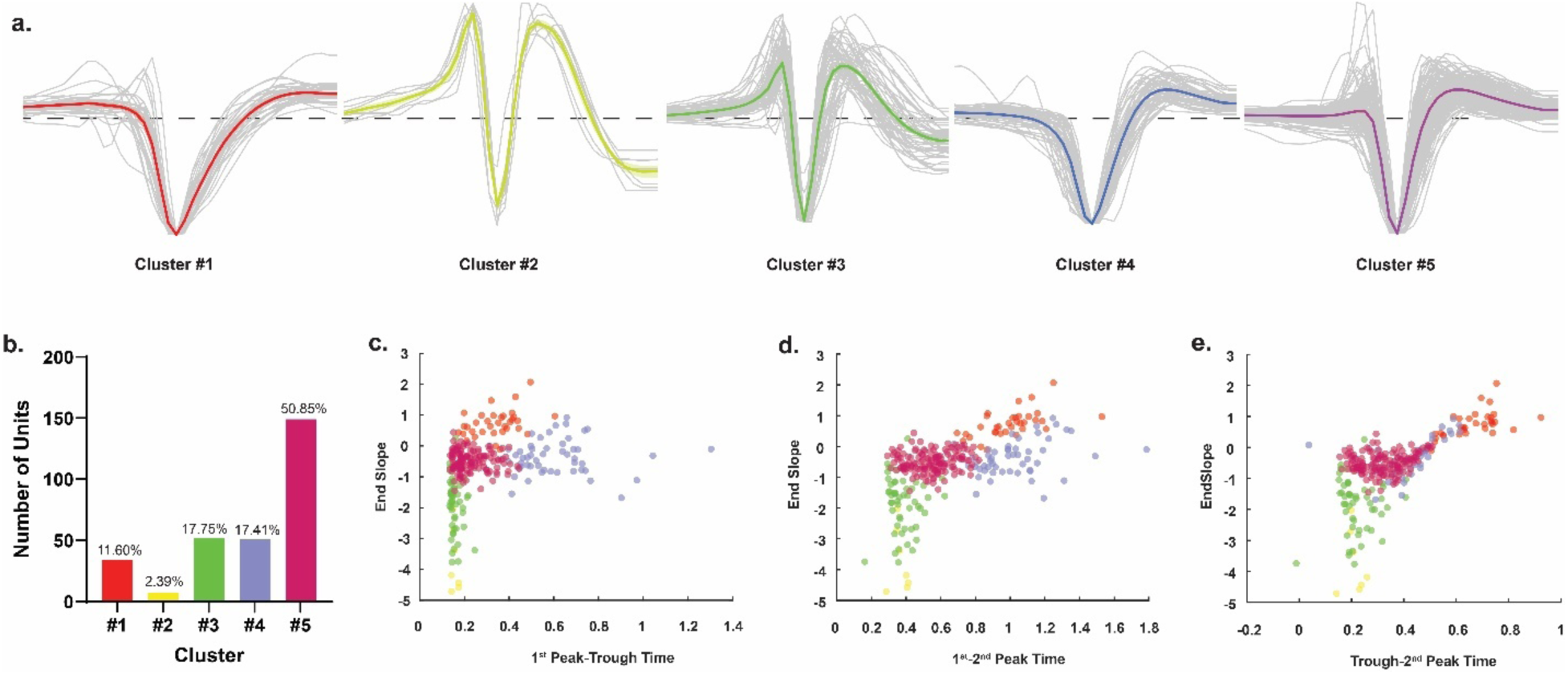
Cluster waveforms, proportions and waveform metric comparison. (a) Individual (*gray*) and mean (*colored*) extracellular spike waveforms relative to baseline (*dashed line*) separated by classification for all SUs. (b) Quantification of total number and percentage of units belonging to each identified cluster. (c-e) Comparison of End Slope and 1^st^-2^nd^ Peak Duration (c), End Slope and Trough-2^nd^ Peak Duration (d), and End Slope and 1^st^ Peak-Trough Ratio (e), for each visually responsive unit with putative clusters indicated by color.

### Classification comparison and validation

To determine the reliability of our semi-automated *k*-means clustering, we used an independent, unbiased method of clustering. Similar to our results above, automated hierarchical clustering also classified the SUs into five distinct clusters (Fig. S3). To further compare classification methods, we generated t-distributed stochastic neighbor embedding plots (t-SNE) plots from both clustering methods (Fig. 4a) [44]. Determining the proportion of each hierarchical cluster comprising each *k*-means cluster revealed that 74% of individual SUs were classified into overlapping clusters (Fig. 4b). In addition, we compared the mean waveforms of each cluster found by the *k*-means and automated hierarchical clustering, which were strikingly similar in all cases (Fig. 4c). Together, these data suggest that both clustering methods broadly generalize to one another.

**Figure 4:**
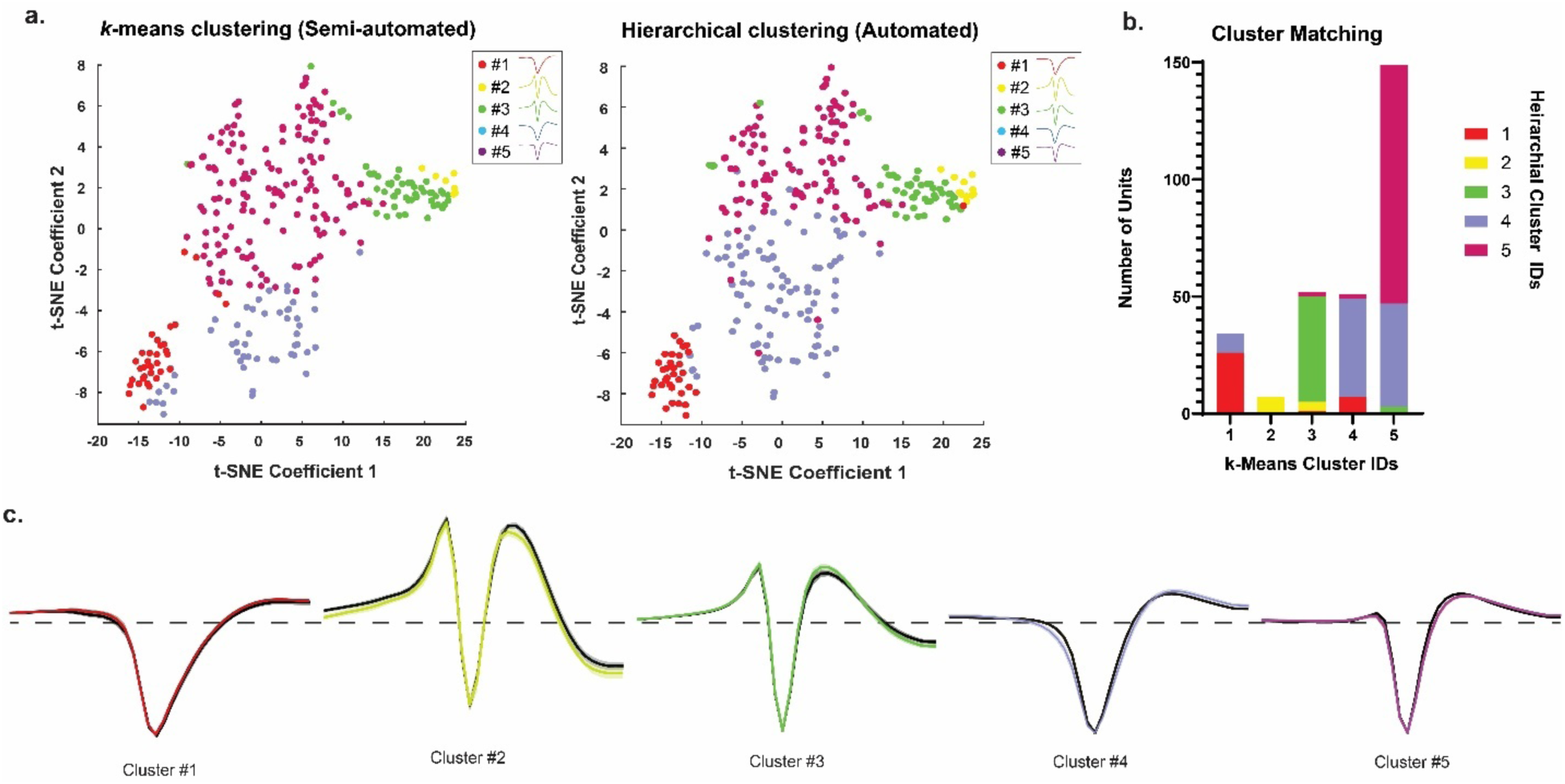
Comparison of cluster composition and waveforms using *k*-means and hierarchical clustering methods. (a) Visualization of units using t-distributed stochastic neighbor embedding (t-SNE). Each data point corresponds to a SU colored by waveform cluster groups from the *k*-means classifier (*left*) and from the hierarchical classifier (*right*). (b) Bar graph showing the total population numbers for each class identified by *k*-means classification, and colored sections within each bar indicating units belonging to the cluster identified by hierarchical classifier. (c) Mean waveforms of clusters identified using *k*-means classifier (*colored*) aligned with mean waveform of matched clusters identified using hierarchical classifier (*black*) relative to baseline (*dashed line*).

From here on, we will use clusters identified by *k*-means clustering for three reasons. First, based on inspection of individual traces and t-SNE plots, hierarchical clustering seemed to include more outliers (Figs. 4A & S3). Second, hierarchical clustering appears to have difficulty segregating Clusters 4 and 5, as evidenced by the inclusion of several waveforms with large amplitude 1^st^ peaks (Fig. S3). Finally, the 2D plots of raw waveform metrics showing SUs suggest better segregation of k-means clusters (Fig. 3c-e) as compared to hierarchical clusters (Fig. S3e-f).

### Firing Statistics of Putative Cell Classes

We next wanted to determine if the putative cell classes identified by waveform-based clustering represented distinct functional populations. We reasoned that each cluster should show different functional characteristics, including firing statistics and visual tuning properties. To begin, we computed four statistics for each neuron in response to visual stimuli: mean firing rate (FR) across all trials, Fano factor (FF), coefficient of variation of the inter-spike interval distribution (CV_ISI_) and burst index (BI). Consistent with each cluster representing distinct physiological populations, we found significant differences in all four measures (p < 0.05, 1-way ANOVA) (Fig. 5a-d). Mean firing rate was highest for class 2 units (8.129 spikes/s) and lowest for class 1 (4.024 spikes/s). We found that the mean rate of units in cluster 3 was significantly different from that of clusters 1 and 5 (P = 0.0284 vs. cluster 1, P = 0.0298 vs. cluster 5, Bonferroni *post hoc* test) (Fig. 5a). For FF, which reflects regularity of firing, class 1 cluster had lowest FF (0.3435±0.05) and cluster 5 had the highest (0.5224±0.02), which were significantly different from one another (P = 0.001). Cluster 4 also had a significantly higher FF from cluster 1 (0.4991±0.04, P = 0.01), indicating that clusters 4 & 5 exhibit less regular firing than cluster 1 (Fig. 5b). Consistent with this, clusters 4 and 5 had elevated CV_ISI_ values (P = 0.0412 cluster 4 v. 3, P = 0.0405 vs. cluster 5 v. 3), a different measure of firing regularity (Fig. 5c). The last metric we examined was BI, which is a measure of how concentrated spikes were when they occurred. We found the most differences between groups based on this index, with cluster 1 having the lowest BI (-0.27±0.098) and cluster 2 having the highest (0.5572±0.12) (Fig. 5d). In addition to these metrics, we compared the mean spike amplitude between groups and again found a segregation between clusters, with cluster 2 having the lowest mean amplitude (-37.04±14.66) and cluster 5 the highest (-252.7±14.80) (Fig. 5e). Importantly, while the ratio of spike amplitude to other peak values was used in our clustering algorithms, absolute amplitude was not. Along with the differences we found in firing statistics between clusters, these data are consistent with the presence of five distinct clusters and support the possibility that each represents a physiologically distinct population in the mouse SC.

**Figure 5:**
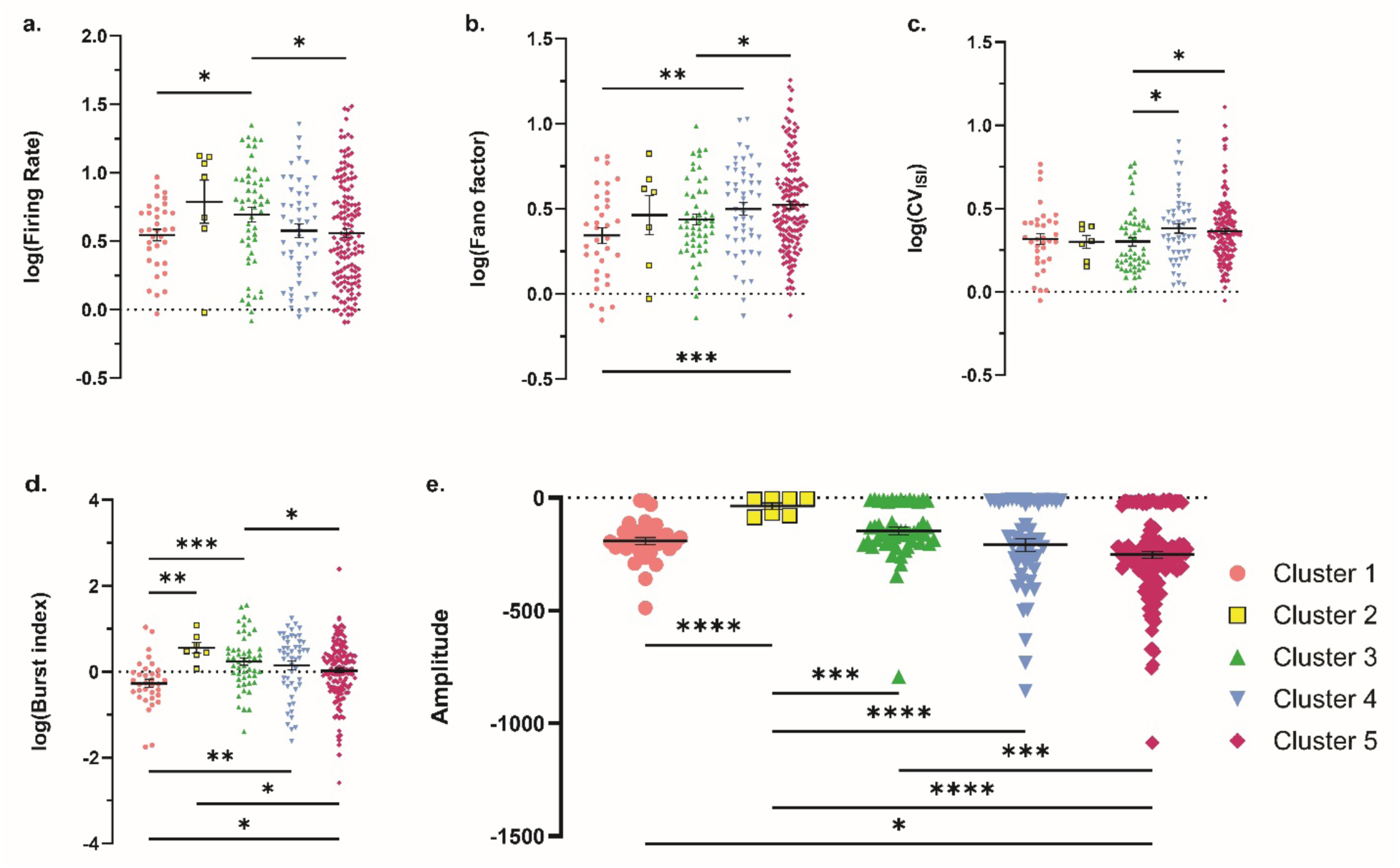
Cluster based analysis of evoked firing statistics. (a) Firing rate: number of times a neuron generates an electrical impulse within a second. (b) Fano factor: variance over mean of spike counts across trials. (c) CV_ISI_: coefficient of variation of the inter-spike interval distribution. (d) Burst index: proportion of inter-spike intervals <5ms over the proportion expected from a Poisson neuron. (e) Amplitude: the difference between the maximum unsigned peak and the baseline (for Cluster 2 it is the difference between the baseline and the height of the 1^st^ Peak).

### Visual tuning properties of putative cell classes

Next, we investigated whether the waveform-based clusters differed in their response to the visual stimulus (Fig. 6). We first determined each neuron’s response to its preferred stimulus and found that units in Cluster 1 had significantly lower evoked firing rates (P = 0.0008 vs. Cluster 3, P = 0.0329 vs. Cluster 4, P = 0.0007 vs. Cluster 5) (Fig. 6a). In contrast, we found no differences in latency to stimulus response (Fig. 6b). Since we used oriented square waves as our stimulus, we determined the gOSI to identify units tuned to orientation. We found significant differences in mean gOSI between Clusters 1 (0.2203±0.03) vs 4 (0.3415±0.03) (P=0.0014) and 1 vs 5 (0.3204±0.15) (P=0.0014), (Fig. 6c). Consistent with this, we found that the proportion of selective units varied (Fig. 6d). We used a cutoff of 0.2 to define a unit as axis selective (AS) and found that ∼50% of SUs in Clusters 1 and 2 were AS, while ∼75% of SUs in Clusters 3, 4, and 5 met this criterion (Fig. 6d). To further understand potential tuning differences between clusters, we examined metrics of orientation tuning for AS units in each cluster. We determine the linearity of spatial summation in SC neuronal responses by calculating an F_1_/F_0_ ratio, the ratio of the response at the drifting frequency to the mean response [40]. Consistent with previous studies [25, 29, 45, 46], most SC neurons exhibited non-linear spatial summation, indicated by an F_1_/F_0_ <1 (242/293; 82.6%). Intriguingly, AS units in Clusters 2 and 3 tended to be more linear than those in Clusters 1, 4, and 5 (mean F_1_/F_0_ [% linear]: Cluster 1 = 0.3522±0.07 [6.67%], Cluster 2 = 1.107±0.35 [66.67%], Cluster 3 = 0.9905±0.08 [57.14%], Cluster 4 = 0.5250±0.07 [13.89%], and Cluster 5 = 0.5779±0.04 [15.53%]---) (Fig. 6e), suggesting clusters may be comprised of distinct populations of AS units. In further support of this, we found that the tuning width (half-width at the half-maximum) of Cluster 4 was minimum (32.25±1.42) and significantly different than Clusters 2, 3, and 5 (P = 0.0034 vs. Cluster 2 (42.10±2.771), P = <0.0001 vs Cluster 3 (41.54±1.577), and P=0.0018 vs Cluster 5 (37.86±0.982)) (Fig. 6f). Taken together with differences found in firing statistics, these data strongly support the presence of five distinct waveform-based cell clusters in the superficial SC.

**Figure 6:**
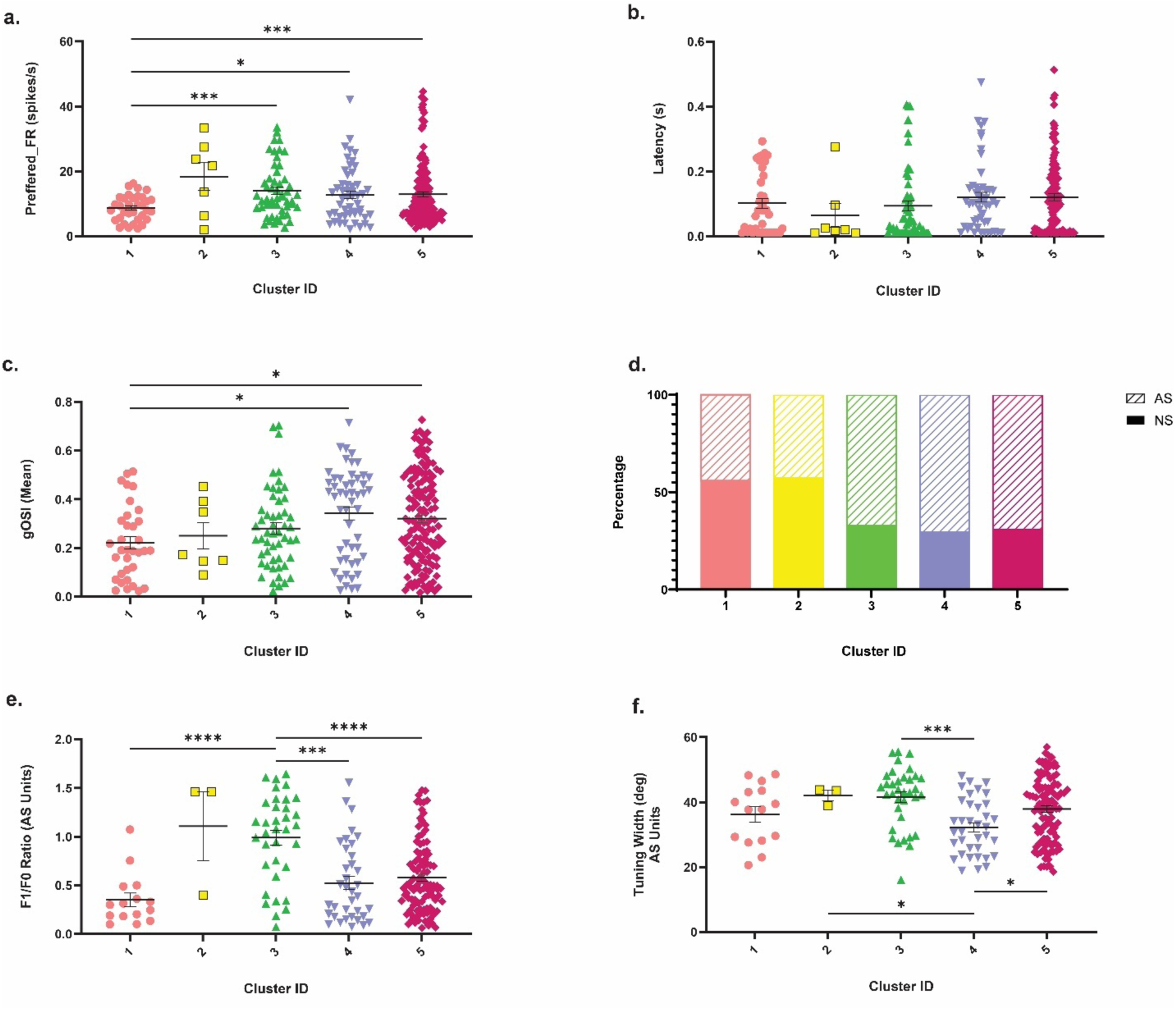
Visual response properties of identified clusters. (a) Preferred FR: number of times a neuron generates an electrical impulse within a second when presented with the stimulus trail. (b) Latency of response after the stimulus is presented. (c) Mean axis selective index (gOSI) across the identified cluster groups. (d) Proportion of orientation selective (AS, striped bars), and nonselective (NS, solid bars) units identified in each identified cluster type. (e) F_1_/F_0_ Ratio for orientation selective units. (f) Tuning width of orientation selective units.

### Alterations of putative cell class composition and function in *Fmr1^-/y^* mice

Based on our previous studies revealing subtle differences in visual function in the SC [32], we wanted to assess if loss of *Fmr1* might impact putative cell classes differently. First, we determined which of the identified SUs were from control or *Fmr1^-/y^* mice. Strikingly, we found that the proportions of identified clusters varied dramatically in the SC of *Fmr1^-/y^* mice. Indeed, we found that no SUs classified as Cluster 2 came from units in *Fmr1^-/y^* mice (Fig. 7a). Additionally, substantially more SUs from *Fmr1^-/y^* mice were classified as Cluster 1 (16.38%, 29/177) compared to those from controls (4.31%, 5/116). Next, we compared firing statistics between genotypes within each cluster. Consistent with proper cluster classification, these were largely consistent between genotypes within a cluster. However, we observed small but significant differences in firing rates between genotypes for Cluster 3 (control: 0.81±0.06, *Fmr1^-/y^*: 0.61±0.07, P = 0.0107) and Cluster 5 (control: 0.67±0.05, *Fmr1^-/y^*: 0.49±0.04, P < 0.0001) (Fig. 7b). We also found decreased FF for Cluster 1 (control: 0.67±0.04, *Fmr1^-/y^*: 0.52±0.05, P = 0.1420) and decreased CV_ISI_ for Cluster 4 (control: 0.55±0.07, *Fmr1^-/y^*: 0.60±0.07, P = 0.6233) (Fig. 7c & d), suggesting less regular firing. Strikingly, we observed dramatic increases in spike amplitude for SUs from *Fmr1^-/y^* mice in all clusters, which reached significance for Clusters 3, 4, and 5 (Cluster 3, P = 0.0005; Cluster 4, P < 0.0001; Cluster 5, P < 0.0001) (Fig. 7e). Taken together, these data suggest that intrinsic firing properties are altered in a subtype-specific manner in the SC of *Fmr1^-/y^* mice.

**Figure 7:**
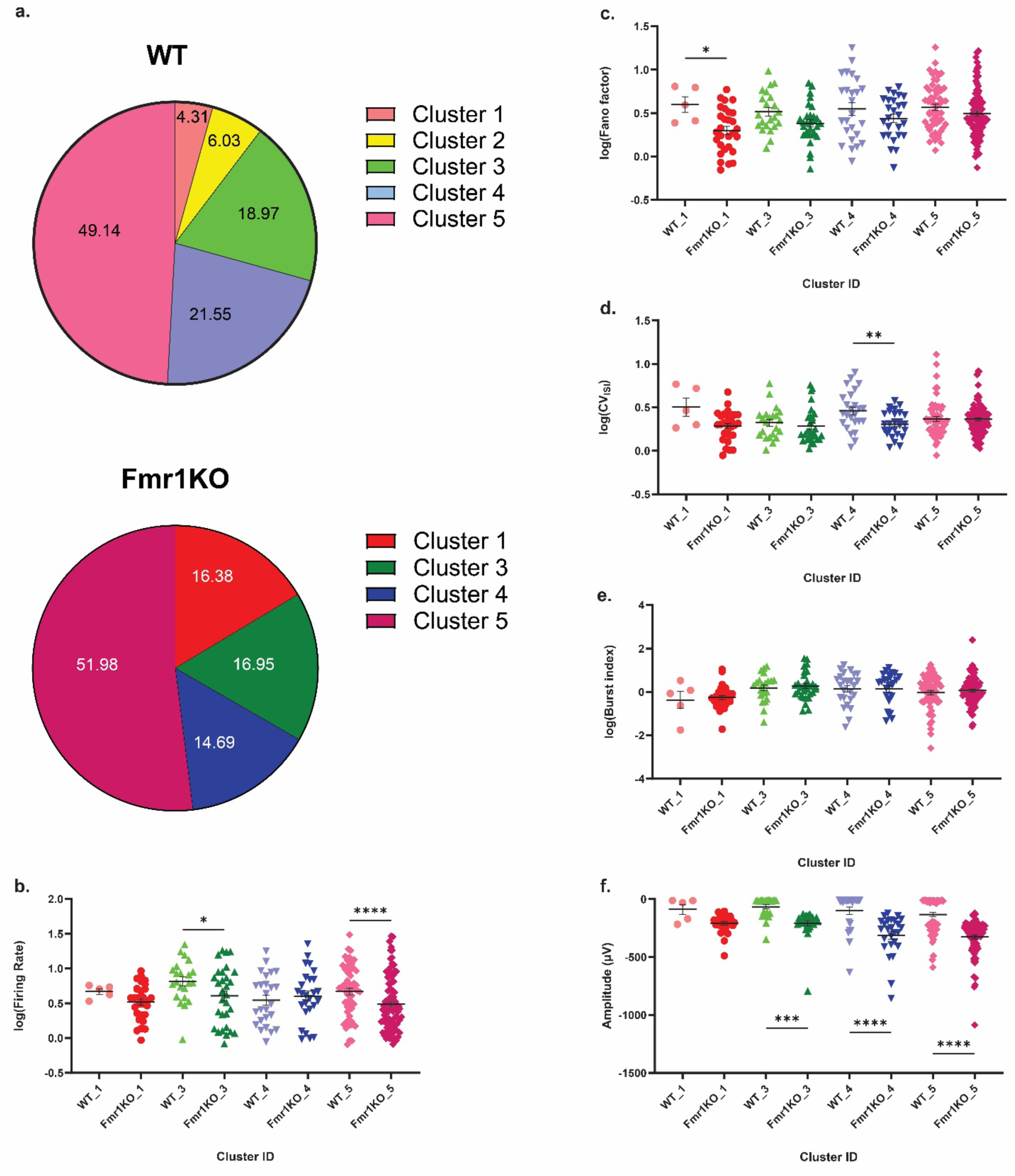
Cluster based analysis of evoked firing statistics in *Fmr1^-/y^* mice. (a) Proportion of SUs belonging to different clusters in control (*top*) and *Fmr1^-/y^*mice (*bottom*). (b-e) Quantification of mean firing rate (b), log transformed Fano factor (c), log transformed CV_ISI_ (d), log transformed burst index (e), and spike amplitude (f) for SUs from control (*lighter shade*) and *Fmr1^-/y^* mice (*darker shade*).

We next asked if visual tuning properties varied within clusters based on genotype. Consistent with our analysis of overall firing rate, we observed a decrease in firing rate to the preferred stimulus for SUs in Cluster 5 of *Fmr1^-/y^* mice compared to controls (control: 11.65±1.33, *Fmr1^-/y^*: 11.95±0.95 P < 0.0044) (Fig. 8a). However, this was not accompanied by an increase in latency to visual response for SUs in any cluster (Fig. 8b). Similarly, we found no differences in mean gOSI nor proportion of AS units between genotypes for any cluster (Fig. 8c & d). Furthermore, no changes in the linearity or sharpness of tuning were observed between genotypes among AS units from any cluster (Fig. 8e & f). Taken together, these data suggest that despite alterations in intrinsic firing properties, loss of FMRP does not impact tuning characteristics of neurons in the SC in a cell class-dependent manner.

**Figure 8:**
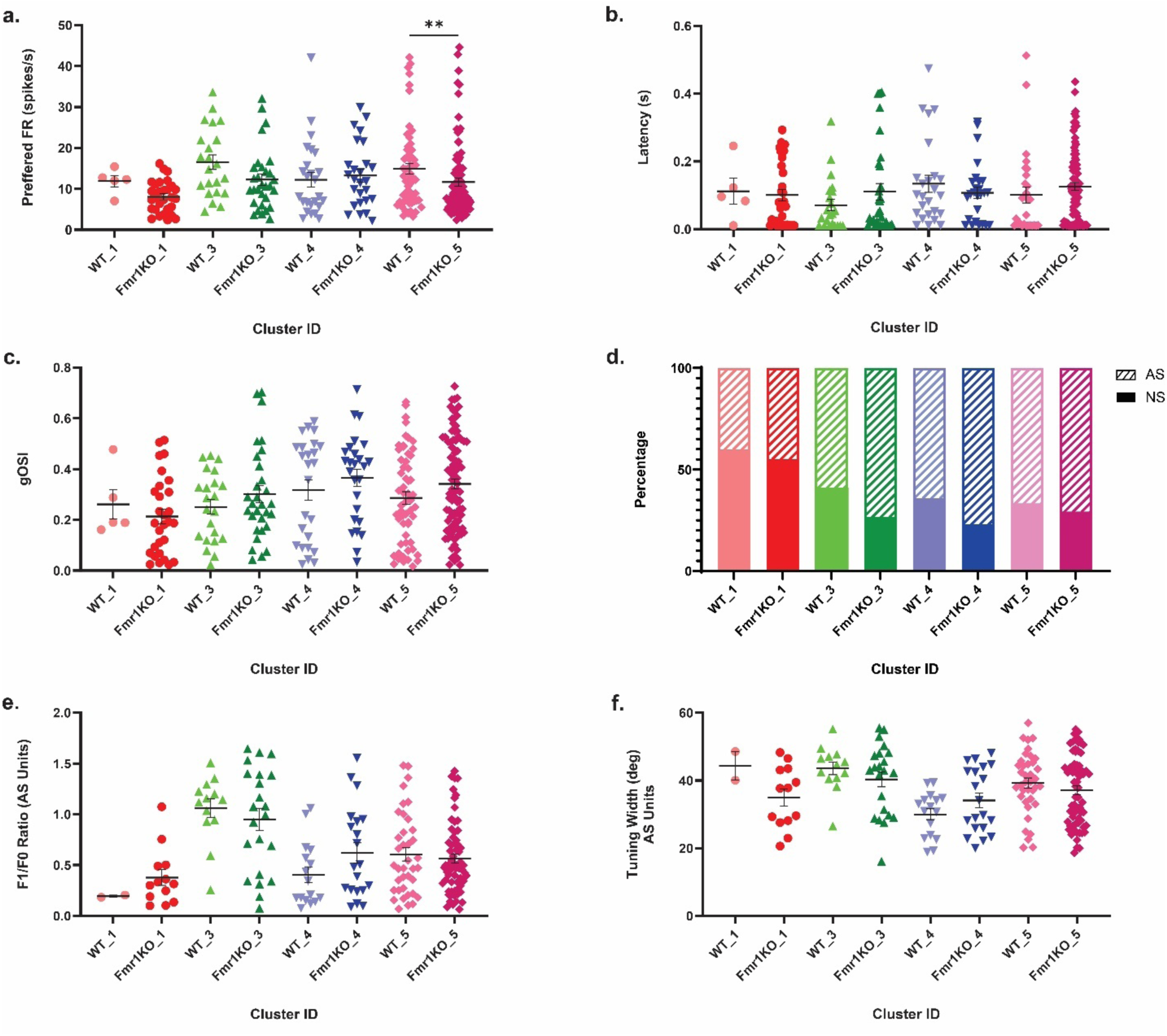
Visual response properties of identified SU clusters in control and *Fmr1^-/y^* mice. (a-c) Quantification of evoked firing rate (a), latency to respond (b), and gOSI (c) for SUs in each cluster separated by genotype. (d) Proportion of axis-selective units in each cluster separated by genotype. (e & f) Quantification of F1/F0 ratio (e) and tuning width (f) of axis-selective units in each cluster separated by genotype.

## Discussion

In this study, we utilized a semi-automated classification system to identify distinct waveform shapes of visually responsive neurons in the superficial layers of the mouse SC. Notably, our classification system included multiple waveform parameters but did not rely on manual decision tree sorting nor computationally intense methods, making it suitable for smaller datasets and likely applicable to multiple brain regions. Furthermore, classification of neuronal subtypes via *k*-means semi-automated clustering was largely replicated using fully automated hierarchical clustering, suggesting the diversity of extracellular signals in the SC is robust. In addition, we observed differences in firing statistics and visual tuning properties between identified subtypes, further supporting the validity of our approach. Intriguingly, our method was also flexible enough to identify within cluster differences between members from different genotypes. We also observed changes in the proportion of neurons in each cluster, suggesting a critical role for FMRP in establishing cell diversity in the SC.

### A robust semi-automated method for extracellular waveform classification

Here, we describe a semi-automated classification method to identify distinct extracellular spike waveform shapes of visually responsive SC neurons. Utilizing six different waveform metrics that describe not only action potential duration during different phases, but also the relative amplitudes of positive and negative deflections, we identified principal components that describe the vast majority of variance to use for *k*-means clustering. After evaluating cluster statistics for a range of *k* values, our data support the presence of five distinct waveform shapes in the mouse SC, which was confirmed using an independent clustering method. This conclusion is further supported by analysis of firing statistics and visual tuning properties, which also revealed differences between groups. Overall, we believe our strategy distinguishes itself from previous methods through its use of a rich set of spike shape features, applicability to smaller datasets, and ease of replicability.

Initial studies easily segregated cortical neurons into two types (broad, regular spiking and narrow, fast spiking) using only two to three metrics, which has been replicable across multiple cortical regions (sensorimotor: [2]; auditory: [13]; somatosensory: [10, 11, 47]; visual: [9, 12]). In contrast, our comparisons of metrics from SC neurons did not yield such dichotomous segregation but was sensitive enough to capture subtle differences between cluster waveforms, suggesting that more than two clusters may be present. Indeed, the same may be true of cortex, as more recent work identified four separable waveforms in the cortex derived from only two metrics [4]. A drawback of that approach is the requirement of a rich dataset of thousands of spikes obtained from multiple cortical regions, which may not be feasible for most studies. Incorporation of additional waveform metrics yielded five separable waveforms from wallaby visual cortex [7]. However, a decision tree classification based on user-defined cutoffs at three progressive waveform metrics was utilized, which may not apply or may need to be determined empirically in different regions.

Methods independent of analyzing raw waveform metrics have also been used to identify diverse subtypes in multiple regions. For example, non-linear dimensionality reduction and graph-based clustering yielded eight discernable waveforms from a reasonably sized data set from macaque motor cortex [5]. Similarly, a semi-supervised deep learning model trained on optogenetically evoked spikes could classify five types of cerebellar neurons with 95% accuracy [8]. However, these approaches are computationally demanding and may require large, well-defined datasets that are not likely accessible to many labs. In contrast, the relatively simple approach utilized here can likely be applied to datasets obtained from different regions and across experimental conditions to identify subtype-specific changes.

### Comparison to previously identified neuronal subtypes in the mouse SC

As the mouse SC has become an increasingly important model to investigate circuit development and sensorimotor transformations, our understanding of neuronal diversity in this critical retinorecipient nucleus has grown. Morphological and intrinsic electrophysiological approaches have identified four to six distinct cell types in the superficial layers of the rodent SC [22-24]. However, functional analyses suggest potentially more diversity within these populations, including axis-selective (both linear and non-linear), direction-selective, motion-selective, looming sensitive, ON, OFF, ON-OFF, suppressed by contrast, locomotion-modulated, and binocularly modulated [25, 29, 45, 46, 48-50]. While some of this selectivity may represent overlapping populations, single cell sequencing studies support the presence of between 10-30 transcriptionally distinct neurons in the superficial mouse SC [27, 28, 51, 52]. Excitingly, an increasing number of lines providing genetic access to SC neurons will help move us toward a unified understanding of cellular diversity by integrating molecular, morphological, and functional data [53].

In this study, we identify five distinct extracellular waveform shapes, but how these relate to those defined above is not clear. To begin, two of the subtypes we identified (Clusters 2 and 3) exhibited a biphasic wave, with a large initial positive deflection prior to its negative trough. Elegant work utilizing high density probes and pharmacologic manipulations suggest these may be signals from incoming retinal or cortical axons [54]. Consistent with this, high density recordings in the hippocampus suggest biphasic spikes are axonal potentials based on analysis of cross-correlation histograms with neighboring excitatory or inhibitory neurons [42]. However, current source density analysis suggests that biphasic spikes can also be attributed to backpropagation in dendrites [43], a phenomenon that has also been reported for widefield neurons in the SC [55]. Thus, the units in Clusters 2 and 3 described here may represent axonal signals, widefield cells, or some other type. Future studies leveraging optogenetic activation of genetically identified neurons are needed to distinguish these possibilities.

Of the three remaining clusters, the most prevalent were SUs in Cluster 5, which had a characteristic narrow spike often associated with inhibitory interneurons [56-58] and comprised ∼50% of the population. This is consistent with studies reporting that 30-60% of neurons in the SC are GABAergic [27, 28, 59, 60]. Notably, while some have reported that GABAergic neurons are predominantly horizontal cells in the SC, others found that stellate and narrow field vertical cells also could be labeled in GAD2-Cre mice [24, 60, 61]. In contrast, neurons in Cluster 1 had characteristic broad spikes associated with excitatory pyramidal neurons [12, 57] in the cortex. Previous studies suggest between 40-60% of neurons in superficial SC are excitatory [28, 62], and labelled neurons had stellate, vertical fusiform, widefield, and multipolar morphologies [63, 64]. Since units in Cluster 1 comprised only ∼12% of the population, they do not likely represent all excitatory neurons in the SC. Units in Cluster 4 had an intermediate waveform shape, which may be consistent with regular spiking excitatory neurons. Together, these populations combine to make up ∼30% of the population, which is consistent with reported densities for excitatory neurons in the superficial-most SC [28].

### Alterations of waveform properties and composition in Fmr1^-/y^ mice

Patients with FXS exhibit a wide range of sensory dysfunctions that manifest in multiple modalities [65]. In the visual domain, patients show impaired magnocellular pathway function, which processes information about stimulus movement, while visual form detection is unaffected [66]. Additionally, recent work revealed that FXS patients performed worse on a visual motion discrimination task than healthy controls [67]. However, the etiology of visual deficits in FXS patients remains unclear, precluding development of effective therapies. Previously, we found that visual neurons in the SC of *Fmr1^-/y^* mice exhibited larger RFs and reduced direction selectivity [32]. These deficits were correlated with disrupted organization of cortical inputs to the SC.

Here, we assessed the impact of FMRP loss on extracellular waveform shape and cellular composition in the SC. Strikingly, we found that units from *Fmr1^-/y^*mice could be classified into only four of the five putative cell types identified by our semi-automated clustering method. As mentioned above, units in Cluster 2 may represent axonal signals. One attractive possibility based on these findings is that these signals correspond specifically to cortical axons, which are disorganized in the SC of *Fmr1^-/y^* mice [32]. Investigations isolating signals from corticocollicular inputs by either electrical or optogenetic activation are needed to confirm this. Intriguingly, we also noted an increase in the proportion of units in Cluster 1 in *Fmr1^-/y^* mice, suggesting altered cellular composition. Consistent with this, we previously found a substantial, but not significant increase in axis selective neurons in the SC of *Fmr1^-/y^* mice [32]. However, alternative methods, such as molecular labeling or transcriptomics are needed to definitively determine if differences in subtype proportions are present in the FXS SC.

Unexpectedly, we found differences in waveform properties between genotypes within individual clusters. Specifically, spike amplitude was almost universally increased across clusters in *Fmr1^-/y^*. While distance from the electrode is likely the biggest determinant for the amplitude of extracellularly recorded spikes [3], FMRP regulates expression of several channels critical for action potential generation and propagation [68-71]. Assuming roughly equal distributions of spike distances for neurons recorded in each population, the increased amplitude observed in *Fmr1^-/y^* mice is likely due to altered expression of one or more of these channels. Modelling studies suggest that factors such as dendritic area, membrane capacitance, and axial resistivity may influence extracellular spike amplitude [72]. Given the well documented increases in dendritic spine density in FXS [73], this may also contribute to the increased spike amplitude observed here. However, whether spine density or dendritic architecture is altered in the SC of *Fmr1^-/y^*mice is not known. In addition, immediately prior neuronal experience can impact amplitude, such that the last spike in a burst has a smaller amplitude than the first [74]. While we did not observe any change in BI between genotypes in any cluster, we did find small decreases in firing rates in two clusters. Future studies are needed to determine the mechanistic cause of increased spike amplitude in a subtype-specific manner.

### Limitations of the study

One limitation of our study is that experiments were conducted in anesthetized mice, where neuronal responses to sensory stimulus may be attenuated compared to those in awake animals [75]. Such a reduction in response strength could result in undetected units and underrepresentation of subtype diversity, especially if specific subtypes are more sensitive to anesthetic modulation. Additionally, we used a single visual stimulus to identify responsive SUs, which may exclude neurons tuned to different aspects of the visual scene. Including a broader range of visual stimuli could help reveal units that respond differently, potentially uncovering novel clusters of functionally distinct neuronal subtypes.

## Acknowledgements

We thank members of the Triplett, Shoykhet and Dean labs for helpful discussions and feedback on the manuscript. This work was supported by NIH grants R01EY025267 and R21EY034660 to J.W.T.

## Notes

### Competing Interest Statement

The authors have declared no competing interest.

## References

1 Connors, B. W., Gutnick, M. J. 1990 Intrinsic firing patterns of diverse neocortical neurons. Trends Neurosci. 13, 99–104. (10.1016/0166-2236(90)90185-d)

2 McCormick, D. A., Connors, B. W., Lighthall, J. W., Prince, D. A. 1985 Comparative electrophysiology of pyramidal and sparsely spiny stellate neurons of the neocortex. J Neurophysiol. 54, 782–806. (10.1152/jn.1985.54.4.782)

3 Henze, D. A., Borhegyi, Z., Csicsvari, J., Mamiya, A., Harris, K. D., Buzsáki, G. 2000 Intracellular features predicted by extracellular recordings in the hippocampus in vivo. J Neurophysiol. 84, 390–400. (10.1152/jn.2000.84.1.390)

4 Trainito, C., von Nicolai, C., Miller, E. K., Siegel, M. 2019 Extracellular Spike Waveform Dissociates Four Functionally Distinct Cell Classes in Primate Cortex. Curr Biol. 29, 2973–2982.e2975. (10.1016/j.cub.2019.07.051)

5 Lee, E. K., Balasubramanian, H., Tsolias, A., Anakwe, S. U., Medalla, M., Shenoy, K. V., Chandrasekaran, C. 2021 Non-linear dimensionality reduction on extracellular waveforms reveals cell type diversity in premotor cortex. Elife. 10, (10.7554/eLife.67490)

6 Sun, S. H., Almasi, A., Yunzab, M., Zehra, S., Hicks, D. G., Kameneva, T., Ibbotson, M. R., Meffin, H. 2021 Analysis of extracellular spike waveforms and associated receptive fields of neurons in cat primary visual cortex. J Physiol. 599, 2211–2238. (10.1113/jp280844)

7 Jung, Y. J., Sun, S. H., Almasi, A., Yunzab, M., Meffin, H., Ibbotson, M. R. 2023 Characterization of extracellular spike waveforms recorded in wallaby primary visual cortex. Frontiers in Neuroscience. Volume 17 - 2023, (10.3389/fnins.2023.1244952)

8 Beau, M., Herzfeld, D. J., Naveros, F., Hemelt, M. E., D’Agostino, F., Oostland, M., Sánchez-López, A., Chung, Y. Y., Maibach, M., Kyranakis, S., et al. 2025 A deep learning strategy to identify cell types across species from high-density extracellular recordings. Cell. 188, 2218–2234.e2222. (10.1016/j.cell.2025.01.041)

9 Connors, B. W., Kriegstein, A. R. 1986 Cellular physiology of the turtle visual cortex: distinctive properties of pyramidal and stellate neurons. J Neurosci. 6, 164–177. (10.1523/jneurosci.06-01-00164.1986)

10 Barthó, P., Hirase, H., Monconduit, L., Zugaro, M., Harris, K. D., Buzsáki, G. 2004 Characterization of neocortical principal cells and interneurons by network interactions and extracellular features. J Neurophysiol. 92, 600–608. (10.1152/jn.01170.2003)

11 Andermann, M. L., Ritt, J., Neimark, M. A., Moore, C. I. 2004 Neural correlates of vibrissa resonance; band-pass and somatotopic representation of high-frequency stimuli. Neuron. 42, 451–463. (10.1016/s0896-6273(04)00198-9)

12 Niell, C. M., Stryker, M. P. 2008 Highly selective receptive fields in mouse visual cortex. J Neurosci. 28, 7520–7536. (10.1523/JNEUROSCI.0623-08.2008)

13 Atencio, C. A., Schreiner, C. E. 2008 Spectrotemporal processing differences between auditory cortical fast-spiking and regular-spiking neurons. J Neurosci. 28, 3897–3910. (10.1523/jneurosci.5366-07.2008)

14 Bruno, J. L., Garrett, A. S., Quintin, E. M., Mazaika, P. K., Reiss, A. L. 2014 Aberrant face and gaze habituation in fragile X syndrome. Am J Psychiatry. 171, 1099–1106.

15 Basso, M. A., Bickford, M. E., Cang, J. 2021 Unraveling circuits of visual perception and cognition through the superior colliculus. Neuron. 109, 918–937. (10.1016/j.neuron.2021.01.013)

16 Hoy, J. L., Farrow, K. 2025 The superior colliculus. Curr Biol. 35, R164–r168. (10.1016/j.cub.2025.01.022)

17 Wurtz, R. H. R., Albano, J. E. J. 1980 Visual-motor function of the primate superior colliculus. Neuroscience. 3, 189–226. (10.1146/annurev.ne.03.030180.001201)

18 Shang, C., Liu, A., Li, D., Xie, Z., Chen, Z., Huang, M., Li, Y., Wang, Y., Shen, W. L., Cao, P. 2019 A subcortical excitatory circuit for sensory-triggered predatory hunting in mice. Nat Neurosci. 22, 909–920. (10.1038/s41593-019-0405-4)

19 Hoy, J. L., Bishop, H. I., Niell, C. M. 2019 Defined cell types in superior colliculus make distinct contributions to prey capture behavior in the mouse. Curr Biol. 29, 4130–4138.e4135. (10.1016/j.cub.2019.10.017)

20 Jun, E. J., Bautista, A. R., Nunez, M. D., Allen, D. C., Tak, J. H., Alvarez, E., Basso, M. A. 2021 Causal role for the primate superior colliculus in the computation of evidence for perceptual decisions. Nat Neurosci. 24, 1121–1131. (10.1038/s41593-021-00878-6)

21 Broersen, R., Thompson, G., Thomas, F., Stuart, G. J. 2025 Binocular processing facilitates escape behavior through multiple pathways to the superior colliculus. Curr Biol. 35, 1242–1257.e1249. (10.1016/j.cub.2025.01.066)

22 Langer, T. P., Lund, R. D. 1974 The upper layers of the superior colliculus of the rat: a Golgi study. J Comp Neurol. 158, 405–436.

23 Edwards, M. D., White, A.-M., Platt, B. 2002 Characterisation of rat superficial superior colliculus neurones: firing properties and sensitivity to GABA. Neuroscience. 110, 93–104. (10.1016/S0306-4522(01)00558-9)

24 Gale, S. D., Murphy, G. J. 2014 Distinct representation and distribution of visual information by specific cell types in mouse superficial superior colliculus. J Neurosci. 34, 13458–13471. (10.1523/JNEUROSCI.2768-14.2014)

25 Wang, L., Sarnaik, R., Rangarajan, K., Liu, X., Cang, J. 2010 Visual receptive field properties of neurons in the superficial superior colliculus of the mouse. J Neurosci. 30, 16573–16584. (10.1523/JNEUROSCI.3305-10.2010)

26 Byun, H., Kwon, S., Ahn, H.-J., Liu, H., Forrest, D., Demb, J. B., Kim, I.-J. 2016 Molecular features distinguish ten neuronal types in the mouse superficial superior colliculus. J Comp Neurol. 524, 2300–2321. (10.1002/cne.23952)

27 Xie, Z., Wang, M., Liu, Z., Shang, C., Zhang, C., Sun, L., Gu, H., Ran, G., Pei, Q., Ma, Q., et al. 2021 Transcriptomic encoding of sensorimotor transformation in the midbrain. Elife. 10, (10.7554/eLife.69825)

28 Liu, Y., Savier, E. L., DePiero, V. J., Chen, C., Schwalbe, D. C., Abraham-Fan, R. J., Chen, H., Campbell, J. N., Cang, J. 2023 Mapping visual functions onto molecular cell types in the mouse superior colliculus. Neuron. (10.1016/j.neuron.2023.03.036)

29 Li, Y. T., Meister, M. 2023 Functional cell types in the mouse superior colliculus. Elife. 12, (10.7554/eLife.82367)

30 Wen, W., Wang, Y., Zhou, J., He, S., Sun, X., Liu, H., Zhao, C., Zhang, P. 2021 Loss and enhancement of layer-selective signals in geniculostriate and corticotectal pathways of adult human amblyopia. Cell Rep. 37, 110117. (10.1016/j.celrep.2021.110117)

31 Kaufmann, W. E., Kidd, S. A., Andrews, H. F., Budimirovic, D. B., Esler, A., Haas-Givler, B., Stackhouse, T., Riley, C., Peacock, G., Sherman, S. L., et al. 2017 Autism Spectrum Disorder in Fragile X Syndrome: Cooccurring Conditions and Current Treatment. Pediatrics. 139, S194–s206. (10.1542/peds.2016-1159F)

32 Kay, R. B., Gabreski, N. A., Triplett, J. W. 2018 Visual subcircuit-specific dysfunction and input-specific mispatterning in the superior colliculus of fragile X mice. J Neurodev Disord. 10, 23. (10.1186/s11689-018-9241-1)

33 Consortium, T. D.-B. F. X. 1994 Fmr1 knockout mice: a model to study fragile X mental retardation. Cell. 78, 23–33.

34 Pretto, D., Yrigollen, C. M., Tang, H.-T., Williamson, J., Espinal, G., Iwahashi, C. K., Durbin-Johnson, B., Hagerman, R. J., Hagerman, P. J., Tassone, F. 2014 Clinical and molecular implications of mosaicism in FMR1 full mutations. Frontiers in Genetics. Volume 5-2014, (10.3389/fgene.2014.00318)

35 Baker, E. K., Arpone, M., Vera, S. A., Bretherton, L., Ure, A., Kraan, C. M., Bui, M., Ling, L., Francis, D., Hunter, M. F., et al. 2019 Intellectual functioning and behavioural features associated with mosaicism in fragile X syndrome. Journal of Neurodevelopmental Disorders. 11, 41. (10.1186/s11689-019-9288-7)

36 Brainard, D. H. 1997 The Psychophysics Toolbox. Spat Vis. 10, 433–436.

37 Pachitariu, M., Sridhar, S., Pennington, J., Stringer, C. 2024 Spike sorting with Kilosort4. Nat Methods. 21, 914–921. (10.1038/s41592-024-02232-7)

38 Rossant, C., Kadir, S. N., Goodman, D. F. M., Schulman, J., Hunter, M. L. D., Saleem, A. B., Grosmark, A., Belluscio, M., Denfield, G. H., Ecker, A. S., et al. 2016 Spike sorting for large, dense electrode arrays. Nat Neurosci. 19, 634–641. (10.1038/nn.4268)

39 Montijn, J. S., Seignette, K., Howlett, M. H., Cazemier, J. L., Kamermans, M., Levelt, C. N., Heimel, J. A. 2021 A parameter-free statistical test for neuronal responsiveness. eLife. 10, e71969.

40 Skottun, B. C., De Valois, R. L., Grosof, D. H., Movshon, J. A., Albrecht, D. G., Bonds, A. B. 1991 Classifying simple and complex cells on the basis of response modulation. Vision Res. 31, 1079–1086.

41 Petersen, P. C., Siegle, J. H., Steinmetz, N. A., Mahallati, S., Buzsáki, G. 2021 CellExplorer: A framework for visualizing and characterizing single neurons. Neuron. 109, 3594–3608.e3592. (10.1016/j.neuron.2021.09.002)

42 Someck, S., Levi, A., Sloin, H. E., Spivak, L., Gattegno, R., Stark, E. 2023 Positive and biphasic extracellular waveforms correspond to return currents and axonal spikes. Commun Biol. 6, 950. (10.1038/s42003-023-05328-6)

43 Jia, X., Siegle, J. H., Bennett, C., Gale, S. D., Denman, D. J., Koch, C., Olsen, S. R. 2019 High-density extracellular probes reveal dendritic backpropagation and facilitate neuron classification. J Neurophysiol. 121, 1831–1847. (10.1152/jn.00680.2018)

44 van der Maaten, L., Hinton, G. 2008 Visualizing data using t-SNE. Journal of Machine Learning Research. 9, 2579–2605.

45 Ito, S., Feldheim, D. A., Litke, A. M. 2017 Segregation of visual response properties in the mouse superior colliculus and their modulation during locomotion. Journal of Neuroscience. 37, 8428–8443. (10.1523/JNEUROSCI.3689-16.2017)

46 Kay, R. B., Triplett, J. W. 2017 Visual neurons in the superior colliculus innervated by Islet2+ or Islet2− retinal ganglion cells display distinct tuning properties. Front Neural Circuits. 11, 73. (10.3389/fncir.2017.00073)

47 Bruno, R. M., Simons, D. J. 2002 Feedforward mechanisms of excitatory and inhibitory cortical receptive fields. J Neurosci. 22, 10966–10975. (10.1523/jneurosci.22-24-10966.2002)

48 Zhao, X., Liu, M., Cang, J. 2014 Visual cortex modulates the magnitude but not the selectivity of looming-evoked responses in the superior colliculus of awake mice. Neuron. 84, 202–213. (10.1016/j.neuron.2014.08.037)

49 Ahmadlou, M., Heimel, J. A. 2015 Preference for concentric orientations in the mouse superior colliculus. Nat Commun. 6, 6773. (10.1038/ncomms7773)

50 Russell, A. L., Dixon, K. G., Triplett, J. W. 2022 Diverse modes of binocular interactions in the mouse superior colliculus. J Neurophysiol. 127, 913–927.

51 Tsai, N. Y., Wang, F., Toma, K., Yin, C., Takatoh, J., Pai, E. L., Wu, K., Matcham, A. C., Yin, L., Dang, E. J., et al. 2022 Trans-Seq maps a selective mammalian retinotectal synapse instructed by Nephronectin. Nat Neurosci. 25, 659–674. (10.1038/s41593-022-01068-8)

52 Cheung, G., Pauler, F. M., Koppensteiner, P., Krausgruber, T., Streicher, C., Schrammel, M., Gutmann-Özgen, N., Ivec, A. E., Bock, C., Shigemoto, R., et al. 2024 Multipotent progenitors instruct ontogeny of the superior colliculus. Neuron. 112, 230–246.e211. (10.1016/j.neuron.2023.11.009)

53 Chen, C., Liu, Y., Cang, J. 2025 Accessing genetically defined cell types in the superior colliculus with transgenic mouse lines. iScience. 28, 112194. (10.1016/j.isci.2025.112194)

54 Sibille, J., Gehr, C., Benichov, J. I., Balasubramanian, H., Teh, K. L., Lupashina, T., Vallentin, D., Kremkow, J. 2022 High-density electrode recordings reveal strong and specific connections between retinal ganglion cells and midbrain neurons. Nat Commun. 13, 5218. (10.1038/s41467-022-32775-2)

55 Gale, S. D., Murphy, G. J. 2016 Active dendritic properties and local inhibitory input enable selectivity for object motion in mouse superior colliculus neurons. J Neurosci. 36, 9111–9123. (10.1523/JNEUROSCI.0645-16.2016)

56 Frank, L. M., Brown, E. N., Wilson, M. A. 2001 A comparison of the firing properties of putative excitatory and inhibitory neurons from CA1 and the entorhinal cortex. J Neurophysiol. 86, 2029–2040. (10.1152/jn.2001.86.4.2029)

57 Insel, N., Barnes, C. A. 2015 Differential Activation of Fast-Spiking and Regular-Firing Neuron Populations During Movement and Reward in the Dorsal Medial Frontal Cortex. Cereb Cortex. 25, 2631–2647. (10.1093/cercor/bhu062)

58 Simons, D. J. 1978 Response properties of vibrissa units in rat SI somatosensory neocortex. J Neurophysiol. 41, 798–820. (10.1152/jn.1978.41.3.798)

59 Mize, R. R. 1988 Immunocytochemical localization of gamma-aminobutyric acid (GABA) in the cat superior colliculus. J Comp Neurol. 276, 169–187. (10.1002/cne.902760203)

60 Whyland, K. L., Slusarczyk, A. S., Bickford, M. E. 2020 GABAergic cell types in the superficial layers of the mouse superior colliculus. J Comp Neurol. 528, 308–320. (10.1002/cne.24754)

61 Endo, T., Yanagawa, Y., Obata, K., Isa, T. 2003 Characteristics of GABAergic neurons in the superficial superior colliculus in mice. Neurosci Lett. 346, 81–84. (10.1016/S0304-3940(03)00570-6)

62 Gehr, C., Sibille, J., Kremkow, J. 2023 Retinal input integration in excitatory and inhibitory neurons in the mouse superior colliculus in vivo. Elife. 12, (10.7554/eLife.88289)

63 Jeon, C. J., Gurski, M. R., Mize, R. R. 1997 Glutamate containing neurons in the cat superior colliculus revealed by immunocytochemistry. Vis Neurosci. 14, 387–393. (10.1017/s0952523800011500)

64 Masterson, S. P., Zhou, N., Akers, B. K., Dang, W., Bickford, M. E. 2019 Ultrastructural and optogenetic dissection of V1 corticotectal terminal synaptic properties. J Comp Neurol. 527, 833–842. (10.1002/cne.24538)

65 Sinclair, D., Oranje, B., Razak, K. A., Siegel, S. J., Schmid, S. 2016 Sensory processing in autism spectrum disorders and Fragile X syndrome-From the clinic to animal models. Neurosci Biobehav Rev. (10.1016/j.neubiorev.2016.05.029)

66 Kogan, C. S., Boutet, I., Cornish, K., Zangenehpour, S., Mullen, K. T., Holden, J. J. A., der Kaloustian, V. M., Andermann, E., Chaudhuri, A. 2004 Differential impact of the FMR1 gene on visual processing in fragile X syndrome. Brain. 127, 591–601. (10.1093/brain/awh069)

67 Goel, A., Cantu, D. A., Guilfoyle, J., Chaudhari, G. R., Newadkar, A., Todisco, B., de Alba, D., Kourdougli, N., Schmitt, L. M., Pedapati, E., et al. 2018 Impaired perceptual learning in a mouse model of fragile X syndrome is mediated by parvalbumin neuron dysfunction and is reversible. Nat Neurosci. 21, 1404–1411. (10.1038/s41593-018-0231-0)

68 Liao, L., Park, S. K., Xu, T., Vanderklish, P., Yates, J. R., 3rd. 2008 Quantitative proteomic analysis of primary neurons reveals diverse changes in synaptic protein content in fmr1 knockout mice. Proc Natl Acad Sci U S A. 105, 15281–15286. (10.1073/pnas.0804678105)

69 Gross, C., Yao, X., Pong, D. L., Jeromin, A., Bassell, G. J. 2011 Fragile X mental retardation protein regulates protein expression and mRNA translation of the potassium channel Kv4.2. J Neurosci. 31, 5693–5698. (10.1523/jneurosci.6661-10.2011)

70 Deng, P. Y., Rotman, Z., Blundon, J. A., Cho, Y., Cui, J., Cavalli, V., Zakharenko, S. S., Klyachko, V. A. 2013 FMRP regulates neurotransmitter release and synaptic information transmission by modulating action potential duration via BK channels. Neuron. 77, 696–711. (10.1016/j.neuron.2012.12.018)

71 Myrick, L. K., Deng, P. Y., Hashimoto, H., Oh, Y. M., Cho, Y., Poidevin, M. J., Suhl, J. A., Visootsak, J., Cavalli, V., Jin, P., et al. 2015 Independent role for presynaptic FMRP revealed by an FMR1 missense mutation associated with intellectual disability and seizures. Proc Natl Acad Sci U S A. 112, 949–956. (10.1073/pnas.1423094112)

72 Pettersen, K. H., Einevoll, G. T. 2008 Amplitude variability and extracellular low-pass filtering of neuronal spikes. Biophys J. 94, 784–802. (10.1529/biophysj.107.111179)

73 Pfeiffer, B. E., Huber, K. M. 2009 The state of synapses in fragile X syndrome. Neuroscientist. 15, 549–567. (10.1177/1073858409333075)

74 Quirk, M. C., Blum, K. I., Wilson, M. A. 2001 Experience-dependent changes in extracellular spike amplitude may reflect regulation of dendritic action potential back-propagation in rat hippocampal pyramidal cells. J Neurosci. 21, 240–248. (10.1523/jneurosci.21-01-00240.2001)

75 De Franceschi, G., Solomon, S. G. 2018 Visual response properties of neurons in the superficial layers of the superior colliculus of awake mouse. Journal of Physiology. 596, 6307–6332. (10.1113/JP276964)

