## Supplemental Figure for "Extracellular spike waveform analysis reveals cell type-specific changes in the superior colliculus of fragile X mice"

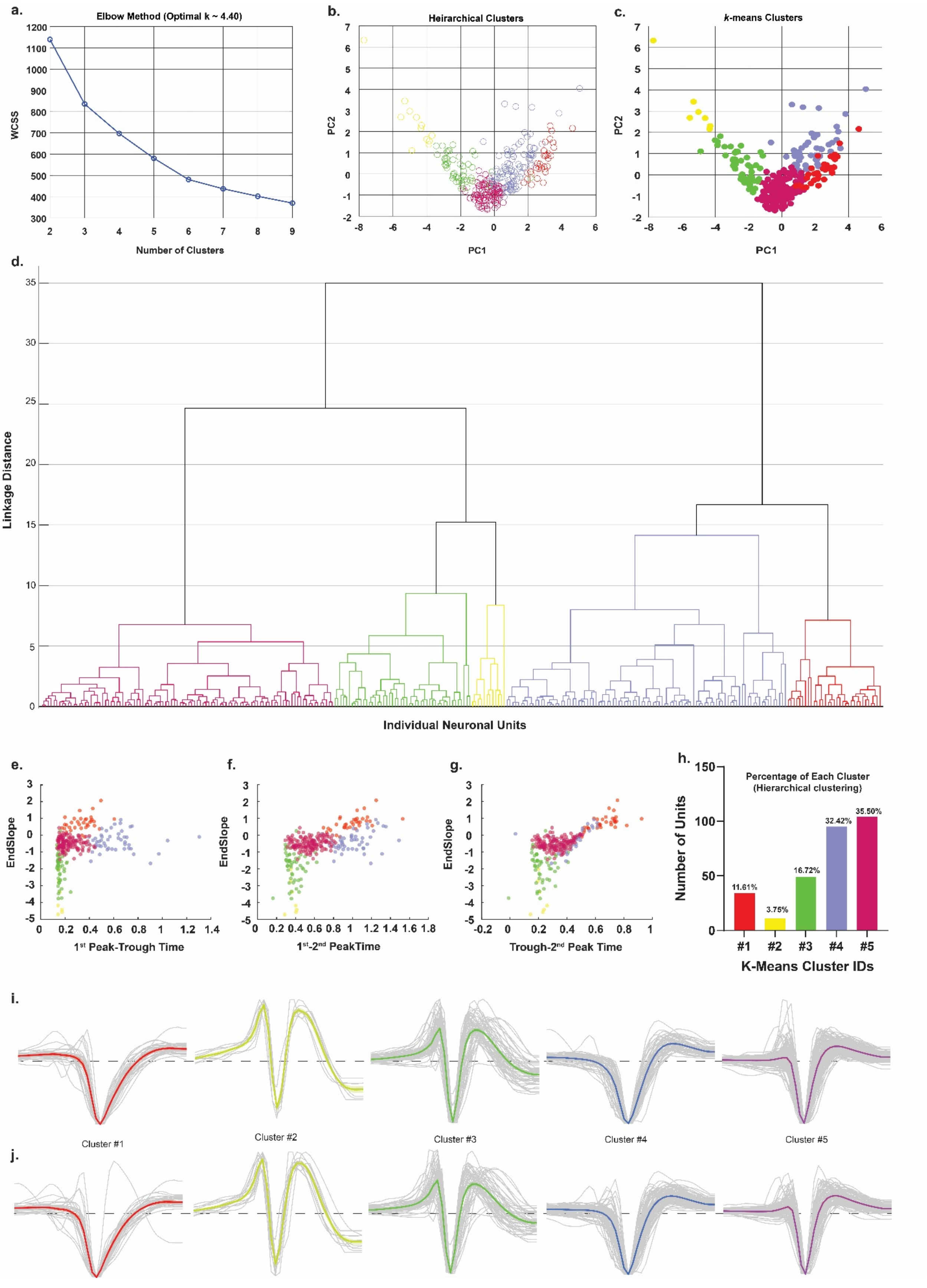
a.
Elbow Method (Optimal k ~ 4.40)
WCSS
Number of Clusters
b.
Heirarchical Clusters
PC2
PC1
c.
k-means Clusters
PC2
PC1
d.
Linkage Distance
Individual Neuronal Units
e.
EndSlope
1st Peak-Trough Time
f.
EndSlope
1st-2nd PeakTime
g.
EndSlope
Trough-2nd Peak Time
h.
Percentage of Each Cluster
(Hierarchical clustering)
Number of Units
11.61%
3.75%
16.72%
32.42%
35.50%
#1
#2
#3
#4
#5
K-Means Cluster IDs
i.
Cluster #1
Cluster #2
Cluster #3
Cluster #4
Cluster #5
j.

**Figure 3 supplementary: Cluster waveforms, proportions and waveform metric comparison.** (a) Plot showing within-cluster sum of squares for the number of clusters using the Hierarchical clustering with optimal cluster number determined using elbow method (*k*=4.4). (b) Two-dimensional PCA projection (PC1 vs PC2) of all units, colored by hierarchical clusters obtained via Ward's linkage and elbow criterion (five clusters). (c) Two-dimensional PCA projection (PC1 vs PC2) of all units belonging to a putative cluster indicated by color, determined by *k*-means clustering algorithm (five clusters). (d) Hierarchical dendrogram (Ward's method) of individual units' assignment to a cluster based on linkage distance with branch colors indicating the five clusters used in (b). (e-g) Relationship between waveform features for each unit belonging to a cluster (indicated by color) obtained via hierarchical clustering. Comparison of End Slope and 1$^{st}$ Peak to Trough Duration (c), End Slope and 1$^{st}$ Peak to 2$^{nd}$ Peak Duration (d), and End Slope and Trough to 2$^{nd}$ Peak Time (e), for each visually responsive unit with putative clusters indicated by color. (h) Quantification of total number and percentage of units belonging to each identified cluster by hierarchical clustering. (i) Individual (*gray*) and mean (*colored*) extracellular spike waveforms relative to baseline (*dashed line*) separated by *k*-means cluster classification for all SUs. (j) Individual (*gray*) and mean (*colored*) extracellular spike waveforms relative to baseline (*dashed line*) separated by hierarchical cluster classification for all SUs.
